# Genomic profiling of 553 uncharacterized neurodevelopment patients reveals a high proportion of recessive pathogenic variant carriers in an outbred population

**DOI:** 10.1101/675868

**Authors:** Youngha Lee, Soojin Park, Jin Sook Lee, Soo Yeon Kim, Jaeso Cho, Yongjin Yoo, Sangmoon Lee, Taekyeong Yoo, Moses Lee, Jieun Seo, Jeongeun Lee, Jana Kneissl, Jean Lee, Hyoungseok Jeon, Eun Young Jeon, Sung Eun Hong, Eunha Kim, Hyuna Kim, Woo Joong Kim, Jon Soo Kim, Jung Min Ko, Anna Cho, Byeong Chan Lim, Won Seop Kim, Murim Choi, Jong-Hee Chae

**Affiliations:** Department of Biomedical Sciences, Seoul National University College of Medicine, Seoul, Republic of Korea; Department of Pediatrics, Seoul National University College of Medicine, Seoul, Republic of Korea; Department of Pediatrics, Gil Medical Center, Gachon University College of Medicine, Incheon, Republic of Korea; Department of Genome Medicine and Science, Gil Medical Center, Gachon University College of Medicine, Incheon, Republic of Korea; Interdisciplinary Program for Bioengineering, Graduate School, Seoul National Universty, Seoul, Republic of Korea; Department of Pediatrics, College of Medicine, Chungbuk National University, Cheongju, Republic of Korea; Department of Pediatrics, Ewha Womans University School of Medicine, Seoul, Republic of Korea

**Keywords:** Whole exome sequencing, Recessive variants, Clinical Neurology, Developmental disorders, Pediatric disease

## Abstract

**Background:** A substantial portion of Mendelian disease patients suffers from genetic variants that are inherited in a recessive manner. A precise understanding of pathogenic recessive variants in a population would assist in pre-screening births of such patients. However, a systematic understanding of the contribution of recessive variants to Mendelian diseases is still lacking.

**Methods:** Genetic diagnosis and variant discovery of 553 undiagnosed Korean patients with complex neurodevelopmental problems (KND for Korean NeuroDevelopmental cohort) were performed using whole exome sequencing of patients and their parents. Pathogenic variants were selected and evaluated based on a comparison to patient symptoms and genetic properties of the variants were analyzed.

**Results:** Disease-causing variants, including newly discovered variants, were identified in in 57.5% of the probands of the KND cohort. Of the 553 patients, 47.4% harbored variants that were previously reported as being pathogenic, and 35.1% of the previous reported pathogenic variants were inherited in a recessive manner. Genes that cause recessive disorders tend to be less constrained by loss-of-function variants and enriched in metabolic and mitochondrial pathways. This observation was applied to an estimation that approximately 1 in 17 healthy Korean individuals carry at least one of these pathogenic variants that develop severe neurodevelopmental problems in a recessive manner. Furthermore, the feasibility of these genes for carrier screening was evaluated.

**Conclusions:** We suggest that the odds are high for healthy individuals carrying a potentially pathogenic variant, and its genetic properties. Our results will serve as a foundation for recessive variant screening to reduce occurrences of rare Mendelian disease patients. Additionally, our results highlight the utility and necessity of whole exome sequencing-based diagnostics for improving patient care in a country with a centralized medical system.

## Background

A large fraction of Mendelian disorders follow a recessive inheritance pattern [1, 2]. The Online Mendelian Inheritance in Men (OMIM) lists 5,317 disorders and 3,077 of these are categorized as recessive (as of April 2019). For more common complex disorders, the contribution of recessive variants to the disease pathogenesis is less than expected [3–6]. For ultra-rare diseases, the contribution of recessive variants in inbred populations, such as Middle-Eastern countries, has been well proven [7–9]. However, the precise contribution of recessive variants to ultra-rare Mendelian disorders in an outbred population is still not well understood.

The inherent complexity of brain developmental processes inevitably leads to patients with diverse neurological problems that frequently challenge conventional diagnostic criteria. Therefore, diagnosis of neurological disorders that affect children is frequently hampered by rare and overlapping clinical features, which makes it difficult for clinicians to readily recognize and properly treat the disease entity. This makes pediatric neurologic patients an impending target for genome-wide genetic studies [10–12]. To facilitate diagnosis and discovery of novel disease pathophysiology, large-scale systematic efforts have been conducted at regional or national scales [13–16]. As many rare neurologic disorders in children follow Mendelian inheritance, disease-causing variant discovery by trio-based whole-exome sequencing (WES) has proved to be the most robust methodology, yielding instant diagnosis rates of 25-41% [10–12, 14, 16].

Notably, the medical system in Korea provides a unique opportunity to conduct a systematic survey of rare disorders and study the contribution of recessive variants at a large scale. With a nation-wide referral system focused on a handful of major tertiary clinical institutions, Seoul National University Children’s Hospital (SNUCH) covers a large portion of the 51-million population, allowing for consistent evaluation and treatment of the patient cohort. For example, we recently reported on genetic analyses of large patient cohorts of Duchenne muscular dystrophy (*n* = 507) and Rett-like syndrome lacking *MECP2* mutations (*n* = 34) [17, 18]. Genetically, ethnic Koreans are a good example of an outbred population in which marriages between relatives and even between individuals possessing the same surnames were prohibited since the 17^th^ century, although the same surname marriage ban was lifted in 2005 [19]. Our study represents the largest of its kind that was conducted at a single clinic, and emphasizes the careful integration of clinical and genetic analyses.

We used WES to analyze a cohort of 553 patients (KND cohort) with severe neurodevelopmental disorders. We characterized the genotype-phenotype relationships of patients whose molecular defects had been identified, and explored the potential association of genes that had not been previously associated with disease. We demonstrate that a high proportion of recessively-inherited variants are associated with patients that have rare neurodevelopmental diseases. Variants that were inherited in a recessive manner were analyzed and their genetic properties were evaluated. Overall, we describe the establishment of a system that efficiently integrates advanced genetic techniques with clinical diagnostic processes to maximize benefits for pediatric patients and their families.

## Materials and Methods

### Subjects

Blood samples were obtained from enrolled patients and their parents, who provided informed consent. WES was performed on 553 patients who visited the SNUCH pediatric neurology clinic from July 2014 to January 2019 and displayed various neurodevelopmental problems of unknown origins, such as demyelinating or hypomyelinating leukodystrophy, hereditary spastic paraplegia, mitochondrial disorders, epileptic encephalopathy, Rett syndrome-like encephalopathy, ataxia, neuropathy, myopathies, and multiple congenital anomalies/dysmorphic syndromes with developmental problems (Table 1). The patients can be categorized into two groups: (i) clinically diagnosable but with substantial genetic heterogeneity (270/553, 48.8%) or (ii) heterogeneous and nonspecific clinical presentations without definite diagnosis (283/553, 51.2%; Additional file: Figure S1). Prior to the WES analysis, thorough clinical and laboratory evaluations and extensive patient examinations have been conducted to identify possible genetic causes. These included genetic tests with candidate gene sequencing, targeted gene sequencing panel, trinucleotide repeat analysis, microarray, metabolic work-up, brain/spine MRI, or muscle biopsy if applicable. All subjects were evaluated by three pediatric neurologists, two pediatric neuroradiologists, and a pathologist.

**Table 1.**
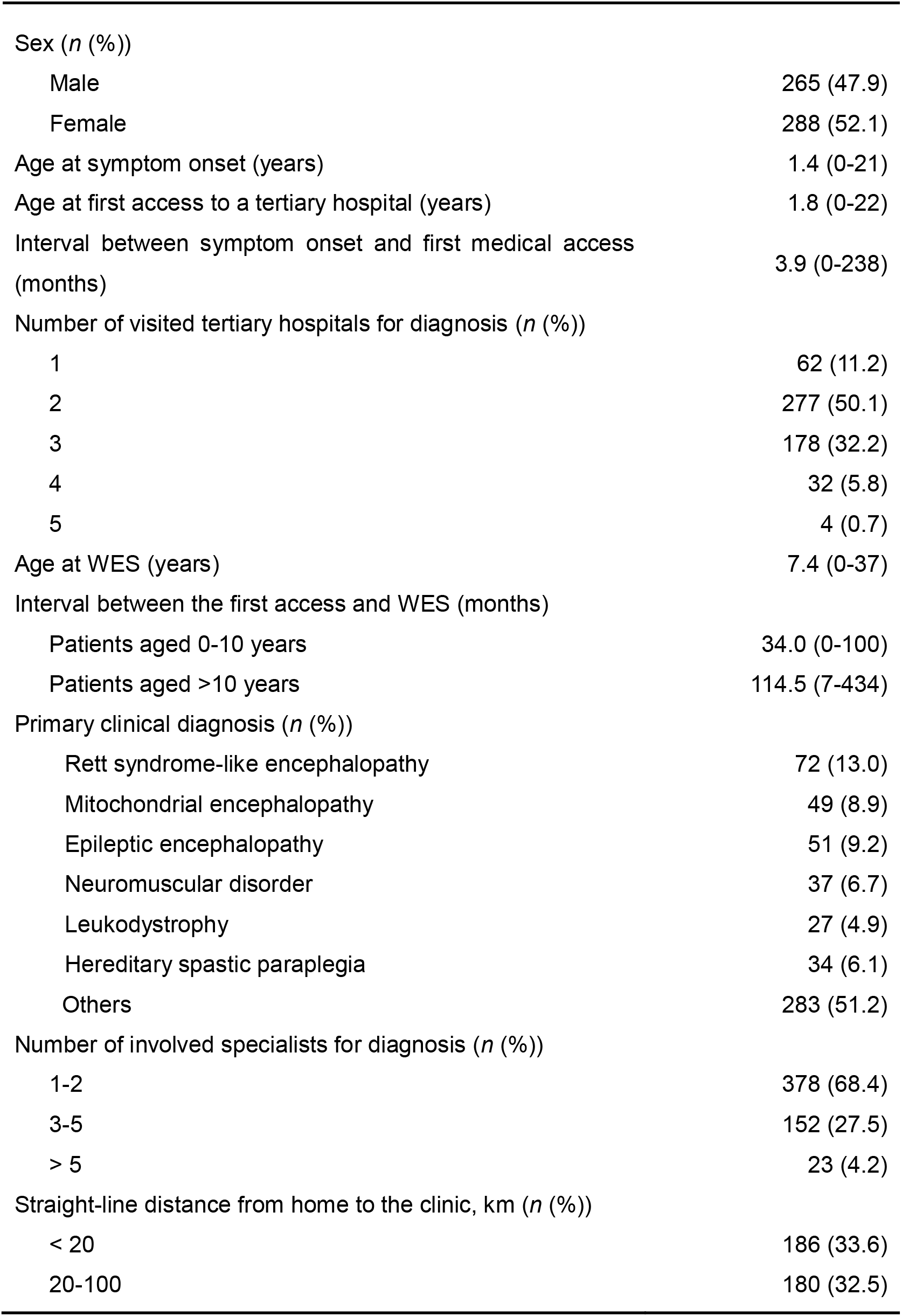
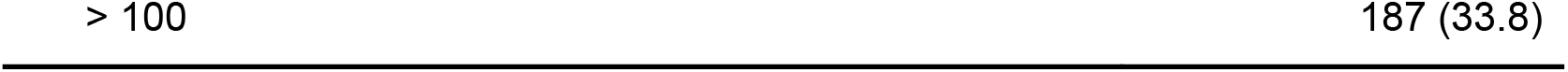
Clinical information of 553 patients.

### Whole Exome Sequencing

WES was performed at Theragen Etex Bio Institute (Suwon, Korea) following the standard protocol and the data were analyzed as described previously [18]. Depending on the genetic analysis result, each patient was categorized as one of the following: category 1: known disease-causing genes were found; category 2: causative gene for other diseases were found; category 3: potentially pathogenic gene, but without prior disease association, was found; category 4: no disease-causing candidates were found; and category 5: known pathogenic copy number variation (CNV) was found.

Our variant assessment procedures were as follows: firstly, patient-specific CNVs were checked and samples with CNVs were classified as category 5. Then, patient-specific nucleotide variants such as *de novo*, compound heterozygous (CH), and rare homozygous (RHo) and hemizygous (RHe) variants were selected by comparing against parents and prioritized based on the inheritance pattern (Fig. 1a). In correlating the patient’s clinical features with genetic variants, if patients carried a known pathogenic variant in OMIM or ClinVar, they were categorized as category 1 or 2, depending on similarity with reported and observed clinical manifestation. For previously unreported variants, if they were never seen in normal individuals (gnomAD [20], Korean Variant Archive (KOVA) [21] and in-house variants), harbored LoF, or were evolutionarily well-conserved at the amino acid level, they were categorized as potentially pathogenic. For CNV calling, the normalized coverage depths of each captured intervals were compared to the depths from related individuals.

**Fig. 1.**
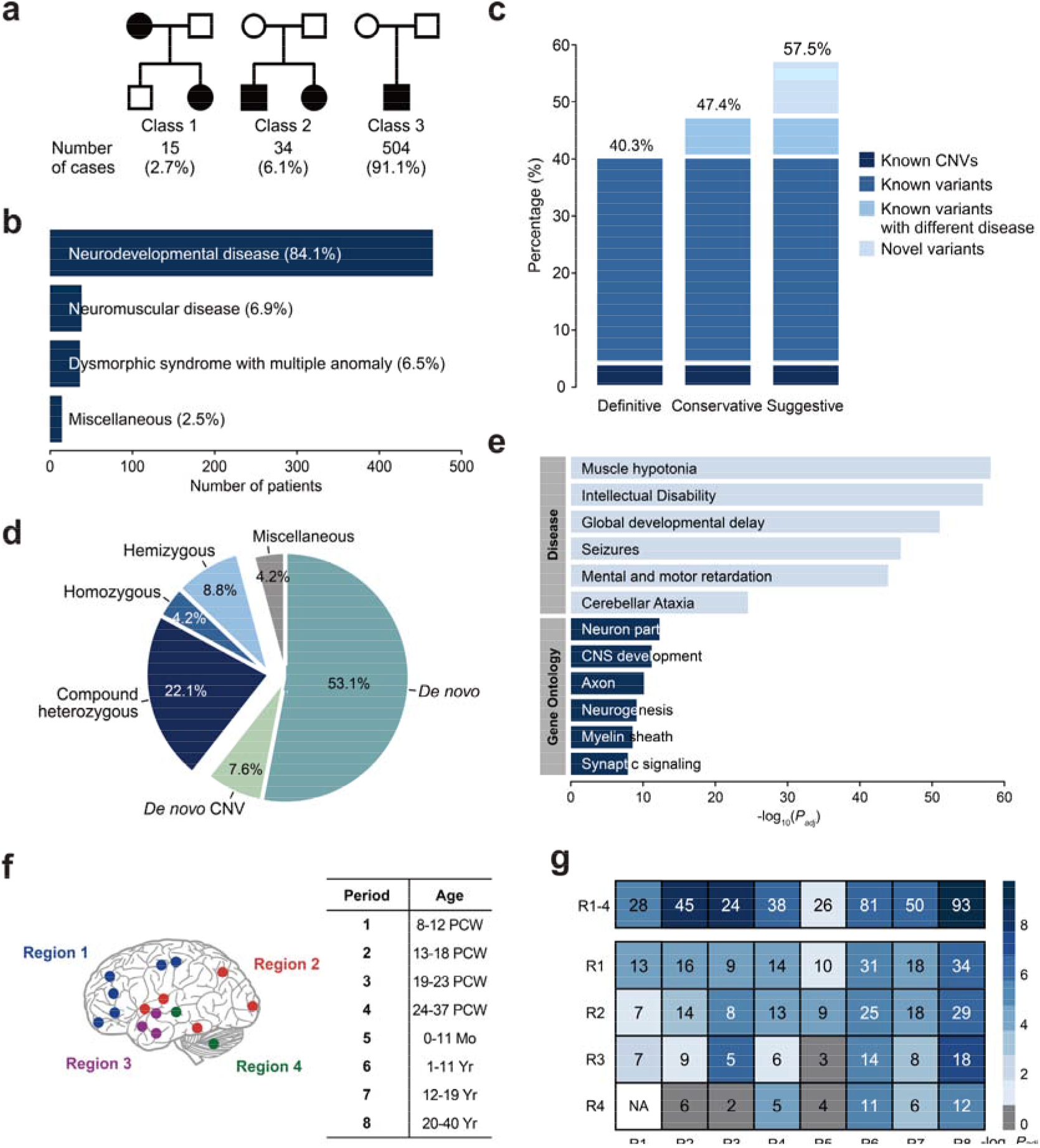
Classification of the KND cohort and results of clinical WES analysis. (a) Subjects by disease inheritance patterns. Class 1: autosomal dominant families; Class 2: families with affected siblings; Class 3: affected individuals with no family history. (b) Major clinical features of the KND cohort. (c) Diagnostic yields of 553 patients with undiagnosed symptoms using WES. (d) Pathogenic variants divided by inheritance patterns. (e) GO and disease enrichment analysis of 164 known genes. (f) Brain anatomical and developmental categorization used for our analysis. Components of each brain region is shown in Additional file: Figure S9. (g) Strength of the co-expression network composed of our known/novel genes compared to random networks as measured by 10^5^ permutations.

### Human Brain Transcriptome Data

The BrainSpan transcriptome database (http://www.brainspan.org) was used to build developing human brain networks [22]. Data from eight post-conceptual weeks to 40 years of age were analyzed. A total of 385 samples were used for the analysis after combining the multiplicates by taking mean values. Probes with TPM (transcripts per million) > 5 in at least one sample were used, yielding 23,943 probes as “brain-expressed transcripts”.

### Brain Transcriptome Network Analysis

Using the above brain-expressed transcripts, we created eight known gene co-expression networks by selecting genes that are highly correlated to our set of genes (*n* = 164; Pearson’s correlation *r* > 0.7), in which their disease associations were previously reported, from each developmental period (Fig. 1f). Then, we asked whether our novel genes can be successfully integrated into the known gene co-expression network. We randomly selected 53 genes (equal to the number of our novel genes) in brain-expressed transcripts and counted how many edges they formed with the known genes. The 10^5^ random gene selections were performed and the number of edges with a known gene was used to construct a distribution. The number of edges from our observed novel genes was evaluated against the random distributions. *P*-values were calculated using z-score, assuming normal distrubutions.

### Recessive variant analysis

Variants were first filtered by gnomAD allele frequency of 0.001 in a heterozygous status and ability to alter protein sequences. Then CH variants were called on a trio setting. If a gene contains more than one filtered variant and each variant was inherited from mother and father separately (for proband), or at least one but not all of the filtered variants of a gene were found from the progeny (for parent), the variants were called as CH. RHo variants were called if filtered variants are inherited in a homozygous manner in autosomes and never seen in gnomAD as homozygous. RHe variants were called if filtered variants are in the X chromosome and never seen in gnomAD as hemizygous or homozygous. Functionality scores were extracted from dbNSFP [23].

## Results

### Diagnostic success rate of WES analyses

The symptoms experienced by KND patients were mostly of pediatric onset (mean 1.4 years of age). Pediatric patients harbored neurodevelopmental problems and were soon referred to tertiary hospitals (mean 1.8 years of age). The majority of the patients visited multiple tertiary hospitals for diagnosis (88.8% visited more than one hospital, mean of 2.3 hospitals), required a mean of 2.3 specialists (31.6% required more than two) and spent a mean of 5.6 years elapsed before WES analysis at SNUCH (Table 1, Additional file: Figure S2). The distribution of straight-line distances from home to the clinic strongly correlates with the original population distribution of Korea, demonstrating that our cohort covers the entire population (Table 1, Additional file: Figure S3).

The majority of the patients is sporadic origin (504/553 = 91.1%; Fig. 1a), making them suitable for trio-based WES analysis. Major clinical feature of the KND cohort was neurodevelopmental disease (84.1%; Fig. 1b). Integrative assessments of genetic variants, their clinical impacts, and patient symptoms allowed us to diagnose 40.3% (223/553) of the cohort with high confidence. The patients included carriers of CNVs (23/553 = 4.2%; 16 heterozygous deletions and 7 duplications; Additional file: Table S1), in which 20 CNVs originated *de novo* (3.6% of the entire cohort), which is slightly lower but comparable to a previous observation [24]. Three inherited pathogenic CNVs were identified: a 165.5 kb deletion in a large family containing multiple affected individuals (Additional file: Figure S4), and 7.2 Mb and 203.2 kb hemizygous duplications that were transmitted from healthy moms to their affected sons (Additional file: Table S1). An additional 7.1% of the cohort (39 patients) harbored previously reported variants that were assumed to be pathogenic but displayed distinct phenotypes, potentially expanding the phenotypic spectrum associated with these genes. For example, two patients that carried a pathogenic heterozygous nonsense or missense variant in *COL1A1*, known to cause osteogenesis imperfecta [25], were initially diagnosed with muscle hypotonia. These two patients did not display skeletal problems, but showed blue sclera [26]. Adding this group to the high confidence group yielded an instant diagnostic rate of 47.4% (“known genes”; Additional file: Table S2). Finally, an additional 10.1% of the cohort (56 patients, 53 genes) harbored variants that are highly likely to be pathogenic but their disease associations are elusive (“novel genes”), yielding a “suggestive” diagnostic yield of 57.5% (Fig. 1c). Among the patients with definite diagnosis, 35.1% are recessive, and 29.9% harbored loss-of-function (LoF) variants (Fig. 1d and Additional file: Figure S5). As expected, the known genes showed strong enrichment in disease categories and gene ontologies such as intellectual disability and central nervous system (CNS) development (Fig. 1e).

### Novel genes display potential enrichment in neuronal differentiation

We assessed whether the 53 novel genes possess a neurologic disease-causing function. The novel gene set was simulated against the BrainSpan data (Materials and Methods) to evaluate if the expression of these novel genes as a group was strongly correlated with known disease-associated genes during brain development. After 10^5^ permutations, we found that the observed involvement of the novel genes was significantly stronger than a randomly selected gene sets across eight developmental windows (Fig. 1f, g; *P*_*adj*_ < 0.05 for all periods). Furthermore, this test was expanded to the four anatomical domains in each period, yielding 32 spatio-temporal windows (Fig. 1g). It is notable that the most highly enriched windows are concentrated in the frontal cortex area (Fig. 1g; R1 × P1-4). These results suggest that expression of the novel genes is concordant with known disease-causing genes in developing brains and this phenomenon is most prominent in the frontal cortex.

### Profile of recessive variants that predispose neurodevelopmental disorders

Using our set of defined pathogenic variants, we explored the genetic properties of these variants that caused disorders in a recessive manner. First, to test if recessive variants (CH, RHo and RHe) are more frequently found in patients as compared to healthy individuals, we counted the number of recessive variants in our cohort and compared these values between patients and healthy parents as controls. Counting all recessive variants from patients and controls, we observed that there is no substantial difference in the number of variants for CH, RHo and RHe (Fig. 2a). Extracting LoF variants, variants in OMIM-listed genes or variants in neurodevelopment-related genes did not reveal any difference in burden (Fig. 2a and Additional file: Figure S6), implying the presence of overwhelming non-pathogenic or non-functional recessive calls in the patients. The majority of genes that were found in our patients with definite diagnosis has been previously documented in OMIM, and has good concordance with previously known recessive or dominant inheritance patterns (Fig. 2b). There were two exceptional cases in which the genes are listed as recessive in OMIM but were dominantly inherited in our patients. First, only the recessive *ACOX1* phenotype has been recognized to date [27], but we are currently working on a report that describes this dominant *ACOX1* variant. Second, a previous report suggested that the *C19orf12* variant has dominant effect, similar to our observations. But this report is not yet listed in OMIM [28]. Next, to test if the genetic properties of recessive variants are different from those of dominant variants, several parameters were compared. Dominant variants (mean allele frequency = 6.2 × 10^−7^) were found less frequently in gnomAD than recessive variants (mean allele frequency = 1.6 × 10^−5^; Mann-Whitney U test *P* = 1.7 × 10^−13^), since most of the dominant variants originated *de novo* whereas recessive variants were inherited from healthy parents (Fig. 2c). Recessive variants were slightly less conserved as compared to the dominant variants, based on PhyloP or amino acid conservation among vertebrate species (Mann-Whitney U test *P* = 0.034 and 0.048, respectively; Fig. 2d). Other functionality test values did not differ significantly between the two groups (CADD *P* = 0.50, GERP *P* = 0.15 and SIFT *P* = 0.17, Mann-Whitney U tests). On the contrary, the genes that contain the recessive variants displayed more lenient constraint as compared to the dominant genes or known haploinsufficiency genes, as documented by observed/expected ratio (o/e) and pLI score in gnomAD [20]. But the recessive genes still display a similar or a slightly more constrained pattern compared to the genes in OMIM (Fig. 2e and Additional file: Figure S7). Functional characterization of recessive neurodevelopmental disease genes revealed an enrichment for genes involved in lipid metabolism and mitochondrial processes (Fig. 2f), in addition to the expected enrichment in CNS development. The relative position of LoF variants in recessive genes demonstrated a similar lenient pattern, more enriched in the C-terminal portion, as compared to the dominant genes (Fig. 2g). There was no significant difference in basic clinical parameters (displayed in Table 1) between the recessive and dominant patient groups (data not shown).

**Fig. 2.**
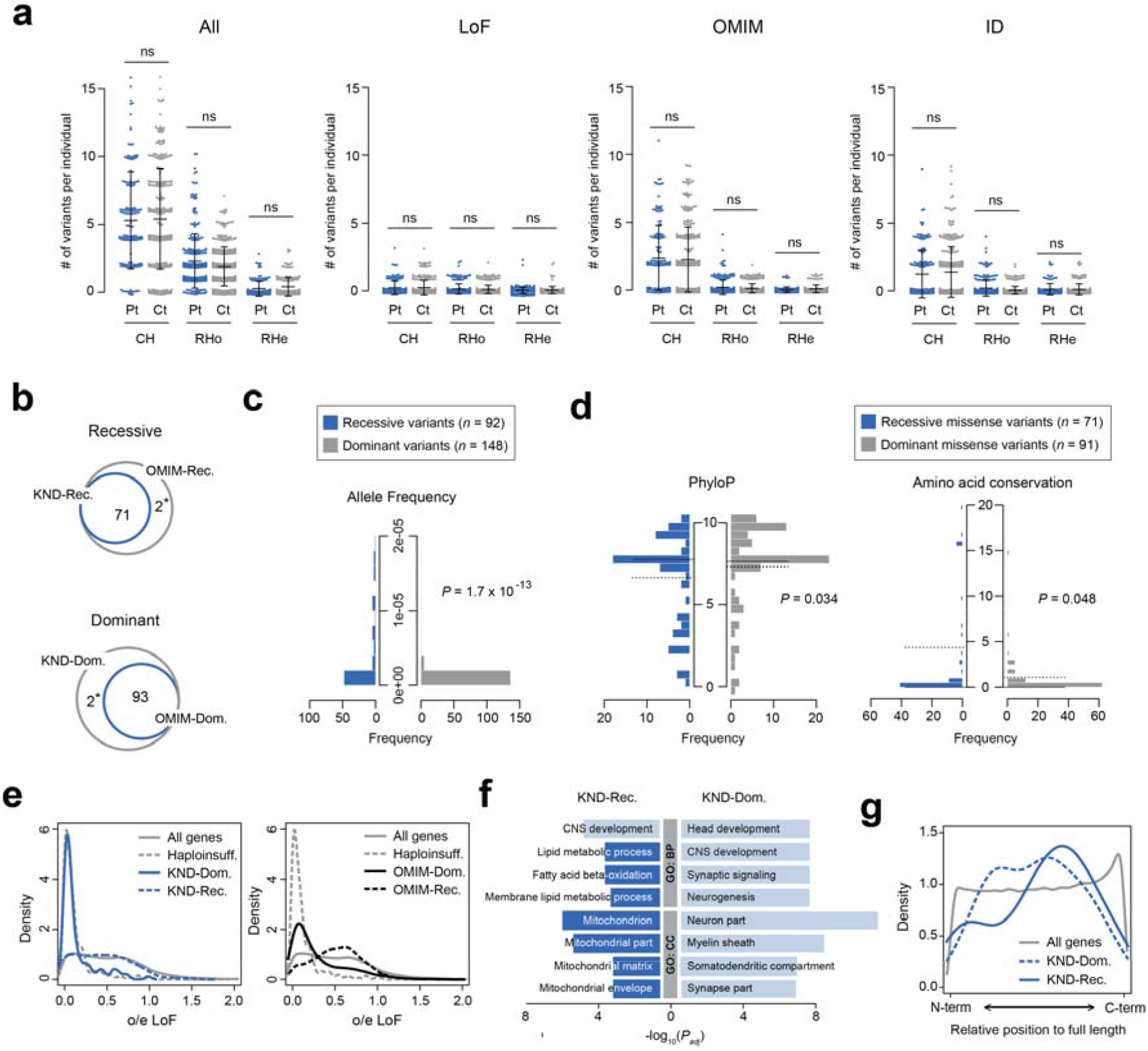
Genetic properties of pathogenic recessive variants. (a) Burden of recessive variants in KND patients (Pt) and their parents as controls (Ct). Recessive variants are divided into compound heterozygous (CH), rare homozygous (RHo) and rare hemizygous (RHe) groups. Numbers found from all variants from all genes (“All”), LoF variants from all genes (“LoF”), all variants from OMIM-listed genes (“OMIM”) and all variants from intellectual disability gene set (“ID”, from DisGeNET) are plotted. Numbers of samples used for each category are as following: patients for CH = 145; controls for CH = 290; patients for RHo = 247; controls for RHo = 341; patients for RHe = 134; controls for RHe = 168. Data are mean ± standard deviation. (b) Venn diagrams displaying high correlations of recessive or dominant inheritance patterns with their known inheritance patterns. The asterisks denote two exceptional cases, *ACOX1* and *C19orf12* (see text). (c) Allele frequency distribution of dominant and recessive variants. (d) PhyloP and amino acid conservation differences between dominant and recessive missense variants. Amino acid conservation is determined by the number of vertebrate species that contain an amino acid that is different from its human orthologous residue. The solid lines denote medians and the dotted lines denote means. (e) Distributions of o/e LoF values for dominant and recessive genes found from KND patients (left) and dominant and recessive genes from OMIM (right) plotted against all genes and known haploinsufficiency genes (*n* = 291) [43]. (f) Functional differences between dominant and recessive genes by GO analysis. Ontologies in dark blue suggest non-neuronal signals specific to the recessive gene group. (g) Relative position of LoF variants in genes. Positions of pathogenic LoF variants in genes from KND patients are plotted against those LoF variant positions from all genes in gnomAD.

### Profile of pathogenic recessive variants carried in healthy individuals

Unlike *de novo* variants that originate largely at random, assembly of recessive variants can be pre-screened and avoided if such variants can be identified in parents. Taking advantage of the extensive coverage of the patient pool maintained by Korea’s centralized medical system and our analysis results of patients with severe neurodevelopmental disorders, it is feasible to infer the probability of recessive variant assembly in Koreans. Several numbers and assumptions are required for this estimation: (i) approximately 400,000 babies are born every year in Korea as of 2016 [29], (ii) approximately 1,000 patients with severe neurodevelopmental disorders newly enroll in our neurodevelopmental disorder clinic every year, (iii) these patients encompass the majority of the Korean population, as exemplified by the sizes of our DMD and Rett cohorts [17, 18] and reflected in the geographical distribution of the KND patients (Table 1 and Additional file: Figure S3), (iv) our result from 553 patients revealed a recessive genetic origin in approximately one-third of the patients (Fig. 1d) and (v) Koreans typically marry an individual with minimal genetic similarity. These observations lead a 1/1,200 incidence rate for developing a severe neurodevelopmental disease in a recessive manner, which can be explained by the existence of one carrier for every 17 healthy individuals (1/1,156; Fig. 3a). One can point out limited evidence for one of our assumptions that we cover the majority of the Korean population. But applying a partial coverage in the estimation will result in increased incidence of neurodevelopment disorder patients and unintentionally inflate the carrier frequency. Therefore, the assumption ensures conservative estimation. Next, we sought to understand the properties of pathogenic recessive variants as compared to the variants found from gnomAD on the same set of 69 genes that contain these variants. As expected, KND recessive variants were found less frequently (Fig. 3b), were strongly conserved during evolution (Fig. 3c), and displayed stronger functionality scores (Fig. 3d) compared to all gnomAD variants found in the same set of genes. Among many variants of obscure functional significances, heterozygous LoF and ClinVar variants in gnomAD can be considered as a first-tier culprit for pathogenic recessive variants. And we observed that the portion that is attributable to LoF and ClinVar variants by healthy carriers was variable among the genes, and this portion is correlated with the o/e LoF value (Pearson’s correlation *r* = 0.33; Fig. 3e).

**Fig. 3.**
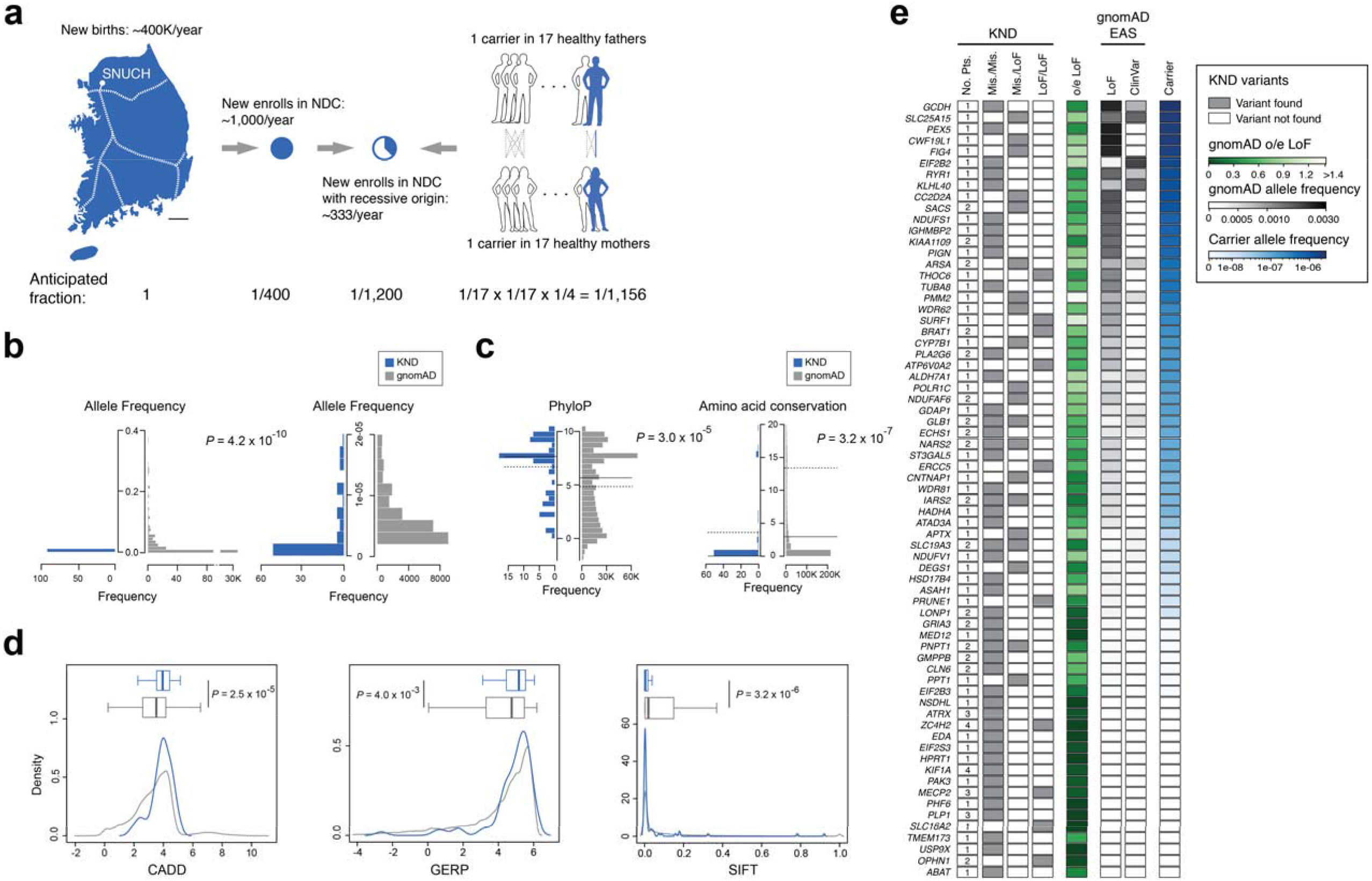
Screen for rare severe neurodevelopmental disorder carriers. (a) A schematic diagram describing processes used to estimate neurodevelopmental disorder carrier frequency in the Korean population. The dotted lines in the map denote the Korea Train Express network, the high-speed railway system of Korea. (b-d) Distribution differences of various parameters between pathogenic recessive variants from KND patients and gnomAD variants from the same genes that were found in KND patients. (b) Allele frequency. The rare frequency portion of the left panel is seperately plotted in the right panel. (c) PhyloP and amino acid conservation. The solid lines denote medians and the dotted lines denote means. (d) CADD, GERP and SIFT. (e) Recessive variants found from KND patients, o/e LoF values, and accumulated frequencies of LoF and ClinVar variants from gnomAD East Asians (EAS) for genes that harbor known pathogenic recessive variants in KND patients. Finally, portion that were attributable to ClinVar or LoF variants for pre-screening parents for each recessive gene are shown.

## Discussion

This study demonstrates the clinical utility of applying WES to children with various and complex neurodevelopmental disorders. We identified genetic causes in 47.4% of the patients and evaluated the characteristics of the variants that caused the disorders in a recessive manner.

Consistent with previous studies, we were able to diagnose approximately half of the KND patients (Fig. 1c) [10–12, 14, 16]. The novel genes formed significantly robust co-expression networks during neurodevelopment processes, which was most prominent in frontal cortex regions (Fig. 1f, g). There was no significant difference in the number of recessive calls between patients and controls, even after stratifying the calls into disease-related gene sets (Fig. 2a and Additional file: Figure S6). This result suggests following: (i) although categorized as “neurodevelopmental”, our patient set is heterogeneous in nature, diluting contribution of single functional entity, (ii) patients carry more non-pathogenic or non-functional recessive calls than expected and we need more power to extract biologically relevant signals. The pathogenic recessive variants that cause rare neurodevelopmental disorders displayed moderately increased allele frequency values and marginally increased evolutionary conservation as compared to dominant variants, suggesting that the qualitative differences between these two groups of variants were not dramatically different (Fig. 2c, d). Compared to the relatively mild differences in variant characteristics, the genes that caused such disorders displayed more discernible differences based on several parameters. First, the recessive genes harbor increased o/e LoF values, implying that there is no constraint applied to the recessive genes, whereas dominant genes in displayed a highly biased pattern toward known haploinsufficiency genes (Fig. 2e). Similarly, the distribution of the relative locations of LoF variants in genes suggested that recessive genes were less constrained compared to dominant genes (Fig. 2g). GO analysis further support that –while the two groups are predominantly composed of neurodevelopment-related genes –recessive genes contain an enriched number of genes that are involved in lipid metabolism and mitochondria function (Fig. 2f). This is a plausible result considering that these pathways are essential for normal brain development [30, 31]. To summarize, these observations suggest that gene properties are stronger determinants of whether the disease adopts a recessive or dominant inheritance pattern than variant properties.

Predicting and avoiding the occurrence of recessive disorders is critical. Carrier estimation has been traditionally performed mostly for single-gene diseases such as β-thalassaemia, Tay-Sachs disease and cystic fibrosis, and efficiently reduced the incidence of these patients [32–34]. However, even after introducing aggressive analysis of genetic disorders using whole exome or whole genome sequencing, precise estimation of the incidence and contribution of rare Mendelian disorders in a recessive manner remains variable for study populations and disorders [2]. For example, analysis of the Deciphering Developmental Disorders (DDD) data suggested only 3.6% of recessive disorders were attributable to patients of European ancestry whereas this value was 30.9% for patients of Pakistani ancestry [4]. Systematic analysis of schizophrenia data did not detect a substantial contribution of recessive variants [5, 6]. These observations differ from ours, where 35.1% of the definitely diagnosed patients emerged in a recessive mode (Fig. 1d), which is in good agreement with previous diagnostic WES studies [10, 35, 36].

Due to a diminishing birth rate and presumably with a burden of having a severely sick child, the majority of our patients lack affected sibling. Therefore, only 11.1% of the patients of recessive origins were from the class 2 pedigree (two siblings being affected, Fig. 1a). Remaining 88.9% of the patients were from the class 3 pedigree and not readily discernable whether their disorders follow dominant or recessive inheritance until WES analysis revealed genetic etiology.

Since the majority of these pathogenic recessive variants were inherited from healthy parents, and ethnic Koreans comprise a relatively isolated population with a centralized medical system, it was feasible to derive an estimate that one out of 17 individuals are healthy carriers of pathogenic recessive variants for severe neurodevelopmental disorders (Fig. 3a). Accumulating this estimated carrier portion across different rare severe disease entities will certainly increase this ratio. The contribution of known LoF and ClinVar variants varies by genes and is positively correlated with o/e LoF values (Fig. 3e), and pathogenic recessive variants display systematic differences that differentiate them from gnomAD variants (Fig. 3b-d). Thus, it would be feasible to predict potential rare recessive variants from genomic data of healthy parents with the help of large patient and control genomic data in the near future.

Our approach expanded the phenotypic spectrum of known genes (39 cases, 7.1%), and suggested novel genes that may allow us to better understand neurodevelopmental disease mechanisms (56 patients, 10.1%). Nevertheless, 42.5% of the cases (235/533) remained undiagnosed even after our WES effort, suggesting a substantial opportunity for further improvement (Fig. 1c). Related to this, a systematic re-analysis effort with additional bioinformatics pipelines increased the diagnostic rate by 4.2% [37]. Also, searching for functional non-coding variants through whole genome sequencing (WGS) and evaluating multiple rare functional variants that may increase disease predisposition may be beneficial [38], although a recent meta-analysis study claimed only a minimal improvement in the WGS diagnostic rate, presumably due to our limited understanding of the function of noncoding variant [39]. An alternative approach would be to integrate genome data with transcriptome data in order to identify functionally cryptic variants that directly influence expression of critical genes [40, 41], although preparing patient-derived tissue still remains as a practical challenge.

Our study also addresses the clinical challenges of an evolving phenotype over time in growing children and how this can be overcome, which facilitates identification of treatable or actionable cases (Table 2; Additional file: Notable vignettes and Figure S8). Our patient cohort included a successful drug repositioning case for a rare neurogenetic disease [42] (Table 2). All of these cases are expected to increase as more genotype-phenotype relationships are discovered and more drugs become available. This study demonstrates that applying WES and subsequent in-depth analysis provides clinical and practical benefits to existing patients and their families and reducing emergence of such patients.

**Table 2.** Notable cases where WES-based analysis conferred correct diagnoses or changed medical treatment strategies.

| Initial clinical problem | Causal gene | Modified clinical interpretation (MIM number) | Significance of WES-based patient evaluation (treatment) | References |
| --- | --- | --- | --- | --- |
| Developmental regression with Rett syndrome-like phenotype | *ST3GAL5* | Salt and pepper developmental regression syndrome (#609056) | Identified the molecular defect and established an accurate diagnosis | [44, 45] |
| Hypotonia and motor delay followed by lower extremity weakness | *DYNC1H1* | Spinal muscular atrophy, lower extremity-predominant 1, AD (#158600) | Diagnosed a case with pleiotropic and evolving symptoms | [46] |
| Early onset hypotonia, sacral mass, congenital heart disease, and facial dysmorphism | *ASAH1* | Farber lipogranulomatosis (#228000) | Corrected a misdiagnosis | [47] |
| Ataxia followed by generalized dystonia | *ANO3* | Expanded spectrum of dystonia 24 (#615034) | Suggested a treatment strategy that resulted in gradual improvement within one year (deep brain stimulation) | [48] |
| Focal lower leg dystonia, dystonic gait | *SLC2A1* | GLUT1 deficiency syndrome 2 (#612126) | Identified disease-specific treatment that resulted in near-elimination of dystonia (ketogenic diet) | [49] |
| Leigh syndrome | *SLC19A3* | Thiamine metabolism dysfunction syndrome 2 (#606152) | Identified disease-specific treatment that resulted in clinical improvements in dystonia, spasticity, and cognitive function (supplements of thiamine and biotin) | [50] |
| Recurrent infections, | *TMEM173* | STING-associated vasculopathy, | Provided a rationale for a new | [42] |
| telangiectatic skin mottling, and brain infarctions |  | infantile-onset (#615934) | treatment strategy that improved the skin lesions (tofacitinib treatment) |  |
| Severe global developmental delay, seizures, and acanthotic skin lesions | *RAB11B* | Neurodevelopmental disorder with ataxic gait, absent speech, and decreased cortical white matter (#617807) | Identified a new disease gene leading to a neurodevelopmental syndrome | [51] |

## Conclusions

We analyzed a comprehensive cohort of rare severe neurodevelopmental disorders in Korea. We genetically diagnosed approximately half of the cohort and discovered novel genetic associations in ~10% of the cohort. Precise etiology for ~40% of the cohort still remains to be elucidated. Extensive analysis of the genetic characteristics of variants and genes that predispose patients to recessive disorders as compared to those of dominant disorders was performed. For this specific set of rare diseases, the properties of genes were a stronger determinant of inheritance pattern as compared to those of variants. Based on these observations, we deduced a ratio of 1/17 for finding a pathogenic recessive variant carrier and suggest several features that predispose variants to reach a pathogenic level. More extensive genome-wide analysis of rare disease patients and healthy controls in a systematic way would provide further insights into the behavior of rare inherited variants that may function as pathogenic recessive variants.

Our study presented a genetic analysis of 553 Korean pediatric patients with unexplained neurodevelopmental problems that revealed various known and novel genetic etiologies. We provided rationales for aggressively extending our system to a wider range of undiagnosed rare disease patients in countries with centralized healthcare like Korea. Through a careful integration of detailed phenotyping, genetic analysis and data sharing, we demonstrate how this approach can facilitate more precise diagnoses and personalized patient care, including pre-screening of rare recessive diseases. Finally, we demonstrated the successful establishment of this approach in Korea, and the necessity of this approach for patients with various undiagnosed neurodevelopmental disorders in countries of similar status.

## Abbreviations

CADD: Combined Annotation Dependent Depletion
CH: Compound heterozygous
CNV: Copy number variation
DDD: Deciphering Developmental Disorders
GERP: Genomic Evolutionary Rate Profiling
gnomAD: Genome Aggregation Database
GO: Gene Ontology
KND: Korean neurodevelopmental disorder
LoF: Loss-of-function
OMIM: Online Mendelian Inheritance in Man
phyloP: phylogenetic P-values
RHe: Rare hemizygous
RHo: Rare homozygous
SIFT: Sorting Intolerant From Tolerant
WES: Whole exome sequencing
WGS: Whole genome sequencing

## Acknowledgements

The authors gratefully acknowledge patients and parents participated in this study.

## Availability of data and materials

Anonymized data not published within this article will be made available by request from any qualified investigator for purposes of replicating procedures and results.

## Authors’ contributions

MC and JHC are responsible for study concept and design, supervised the study and obtained funding. YL, SP, JC, YY, SL, TY, ML, JS, Jeongeun Lee, JK, EYJ, EK, and MC analyzed genome data. JSL, SYK, JC, HK, WJK, JSK, JMK, AC, BCL, WSK, and JHC provided clinical data. YL, SP, JSL, SYK, JC, MC, and JHC combined genetic and clinical data. YL, SP, JC, Jeongeun Lee, JK, Jean Lee, HJ, EYJ, SEH, and MC performed genetic and statistical evaluation of the cohort. YL, SP, JSL, SYK, MC, and JHC drafted the manuscript. All authors reviewed the manuscript for important intellectual content.

## Ethics approval and consent to participate

This study was approved by the Seoul National University Hospital Institutional Review Board (No. 1406-081-588).

## Consent for publication

Not applicable

