## Supplementary material for "Genomic profiling of 553 uncharacterized neurodevelopment patients reveals a high proportion of recessive pathogenic variant carriers in an outbred population"

< Additional Information >

**<Contents>**

**Notable vignettes** 3

**Supplementary Figures**

**Figure S1.** Patients categorized by clinical diagnosis (n = 553 patients). 6

**Figure S2.** The age distribution of all patients when their symptom began and the distance between age of onset to WES analysis (*n* = 553 patients). 7

**Figure S3.** Straight-line distances from SNUCH. 8

**Figure S4.** Identification of an inherited deletion at 2p22.3 in the family with hereditary spastic paraplegia (HSP). 9

**Figure S5.** Pathogenic variants categorized by function (n = 262 variants). 11

**Figure S6.** No difference in the number of recessive variants for neurodevelopment-related gene sets between patients and controls. 12

**Figure S7.** Distribution of pLI values 13

**Figure S8.** Presentation and validation of cases that WES-based analysis altered clinical courses. 14

**Figure S9.** Anatomical components of each brain regions that were used for novel gene network analysis 16

**Supplementary Tables**

**Table S1.** List of copy number variations discovered in this study. 17

**Table S2.** List of known neurologic disorder associated genes. 20

***Notable vignettes***

WES-based diagnoses of 553 patients with neuro-developmental problems generated cases that were clinically meaningful for a number of reasons (Table 2).

*ST3GAL5 case: genetic elucidation of a case of compound symptoms* [1]

Two female siblings presented with a Rett syndrome-like phenotype, such as psychomotor retardation/regression, delayed speech, hand stereotypies with a loss of purposeful hand movements, and choreoathetosis. Genetic tests of *MECP2/CDKL5/FOXG1* and chromosomal microarray found no plausible candidates. GM3 synthase deficiency with *ST3GAL5* compound heterozygous variants was diagnosed through WES, which was confirmed by liquid chromatography-mass spectrometry analysis [1, 2]. WES ended three years of an undiagnosed period and broadened the phenotypic spectrum of this rare neurometabolic disorder.

*DYNC1H1 case: a case of evolving symptoms*

An 8-month-old girl from healthy parents presented with global developmental delay and generalized hypotonia. Laboratory findings, MRIs, electromyography/nerve conduction studies, and genetic tests including *SMN1*, failed to identify her etiology. Although her cognitive function regression improved rapidly to near normal, a suspicious paraplegic gait developed from three years of age. WES revealed a novel *de novo* missense variant in *DYNC1H1*, encoding a subunit of the cytoplasmic dynein complex (Fig. S8a-d). This observation led us to conclude that the patient started displaying global DD, but was eventually diagnosed with SMALED (spinal muscular atrophy, lower extremity-predominant), mimicking hereditary spastic paraplegia. This case poses a rare example, along with recent recognition that *DYNC1H1* carriers display complex HSP [3], which demonstrates the clinical utility for pediatric cases with phenotypic pleiotropy and symptom evolution.

*ASAH1 case: a case with corrected diagnosis* [4]

An 11-month-old girl from healthy parents was referred to our hospital for developmental delay with hypotonia, facial dysmorphism, congenital heart defects, and a sacral mass since birth. Karyotyping, metabolic screening, and chromosomal microarray revealed no abnormalities. Excision of the sacral mass led to pathologic diagnosis of epithelioid hemangioendothelioma, which required repeated operations and chemotherapy. However, joint contracture and multiple subcutaneous nodules appeared from 18 months of age. WES revealed compound heterozygous variants in *ASAH1*, associated with Farber disease. After the establishment of the correct molecular diagnosis at the age of four, additional electron microscopic findings of previously excised masses confirmed the pathognomonic findings [4].

*SLC2A1 case: a case where an effective treatment was given*

A 16-year-old boy from an otherwise a healthy family displayed an abnormal gait from around eight years of age that later developed into falls with dystonia, notably in the lower extremities, which was provoked mainly by exercise or stress. His motor development was delayed, although he had normal head circumference and cognition. WES revealed a *de novo* LoF variant in *SLC2A1*, encoding a glucose transporter (GLUT1) [5] (Fig. S8e-g). We also retrospectively noticed that he had developed episodes of staring spells from an age of four years, suggesting absence seizures, which disappeared after three years of antiepileptic medication. Cerebrospinal fluid analysis confirmed a low glucose level (37 mg/dL, 36% of serum glucose, normal range > 40%). Based on this observation, a ketogenic diet was applied and completely changed the quality of his life, with a resulting near-absence of dystonia.

*RAB11B case: a case of newly identified disease through international data sharing* [6]

A 2-year-old boy was referred for evaluation of global developmental delay. He showed microcephaly and severe cognitive impairment with epilepsy but without evidence of regression. Notably, abnormal acanthotic skin lesions progressed from the face to the whole body. We identified a *de novo* variant in *RAB11B* and submitted this to GeneMatcher [7]. Soon, four cases with a similar neurodevelopmental phenotype were matched and we demonstrated that this gene causes intellectual disability and a distinctive brain phenotype [6].

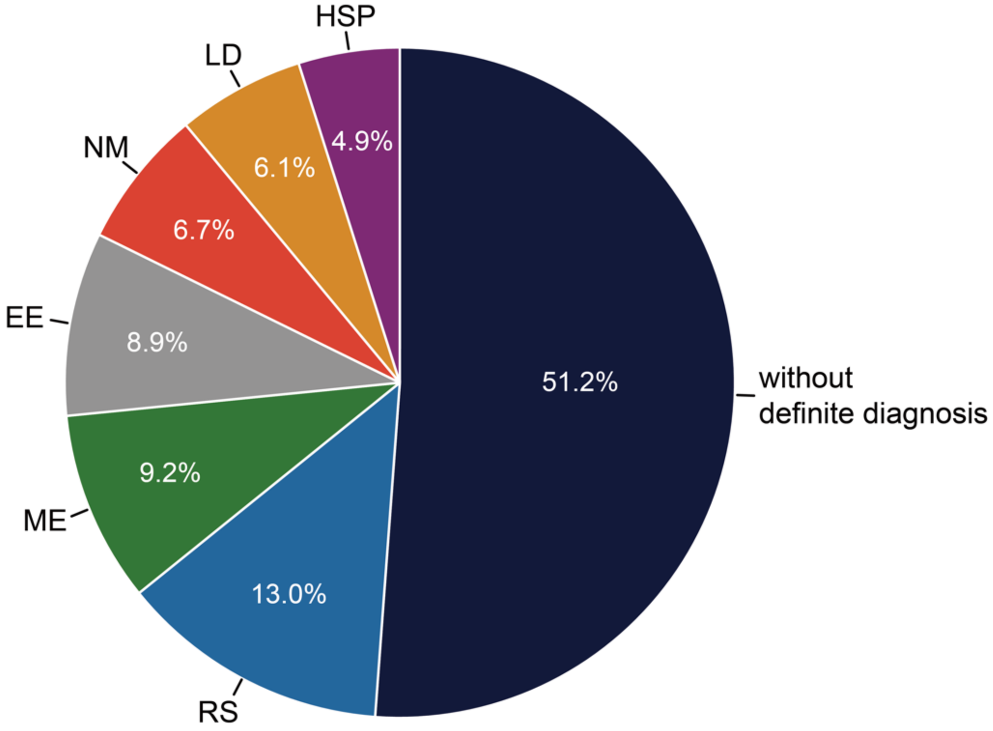


**Figure S1. Patients categorized by clinical diagnosis (*n* = 553 patients).** RS, Rett syndrome-like encephalopathy; ME, Mitochondrial encephalopathy; EE, Epileptic encephalopathy; NM, Neuromuscular disorder; LD, Leukodystrophy; HSP, Hereditary spastic paraplegia

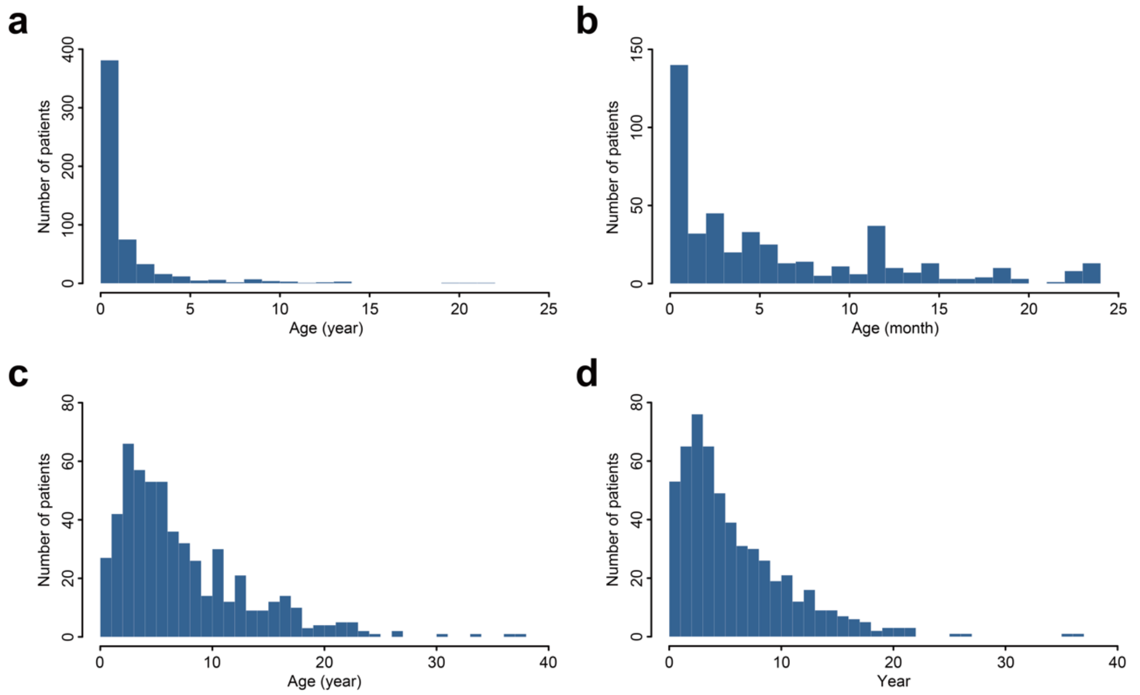


**Figure S2. The age distribution of all patients when their symptom began and the time differences between age of onset to WES analysis (*n* = 553 patients).** (a) The age of onset of all patients (year). (b) The age of onset of patients whose symptoms began before 24 months. (c) The Age distribution of all patients when the WES analysis was performed. (d) Distribution of time differences between patient's age of onset and the time of WES analysis.

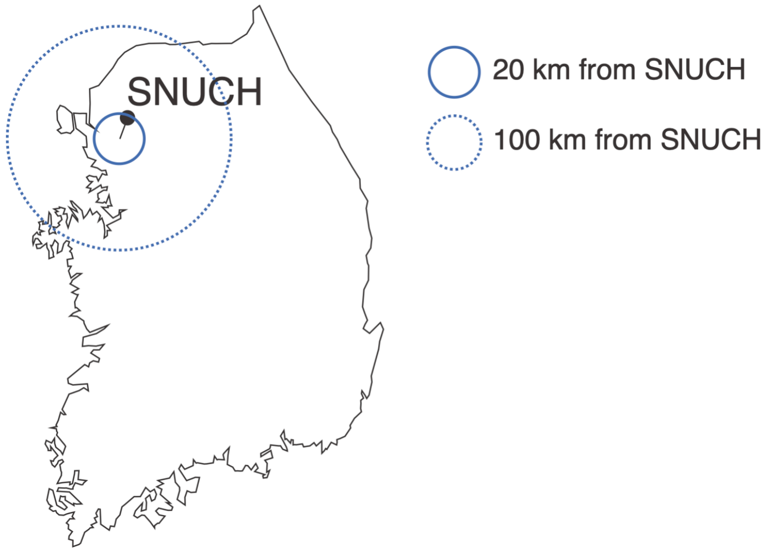


**Figure S3. Straight-line distances from SNUCH.** The 20 km radius circle includes most of Seoul, where about a quarter of entire population resides. The 100 km radius circle encompasses most of the Kyeonggi province, where another quarter of entire population resides.

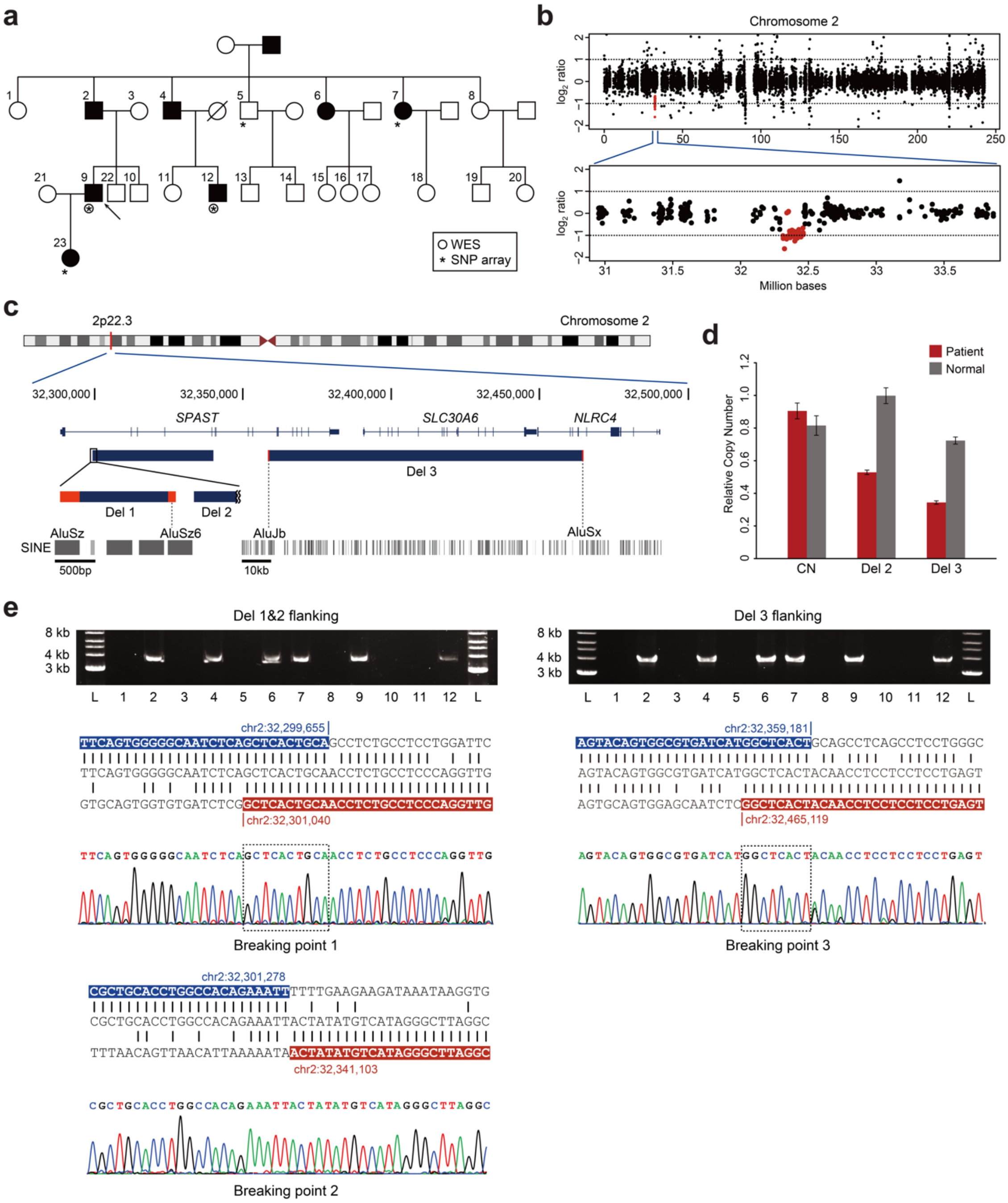


**Figure S4. Identification of an inherited deletion at 2p22.3 in the family with hereditary spastic paraplegia (HSP).** (a) Pedigree of an HSP family with 8 affected individuals across four generations. (b) WES-based log2-based copy-number values of subject HSP-9 compared with an unrelated normal subject on chromosome 2 showing the presence of heterozygous microdeletion. Captured intervals of copy-number loss are indicated by red dots. (c) Enlarged view of deletion-bearing region at 2p22.3

encompassing *SPAST*. Blue solid bars represent the deleted intervals. Alu repeat elements at the deletion breakpoints are indicated by red solid bars and aligned with the UCSC Genome Browser RepeatMasker, represented by dotted lines. (d) Validation of the deleted regions by quantitative PCR of genomic DNA. The red bars represent an average copy-number of five patients and the gray bars represent an average copy-number of five normal individuals from the family. Error bars represent standard error. CN: copy-neutral region. (e) DNA sequence analysis of the deleted regions. The DNA fragments containing the deletion was amplified by the deletion-specific primer pairs. The deletion-specific PCR products of 3.8 kb (Del 1 & 2) and 4.0 kb (Del 3) are observed in each affected individual. Reference sequences surrounding the breaking points are indicated in blue and red color.

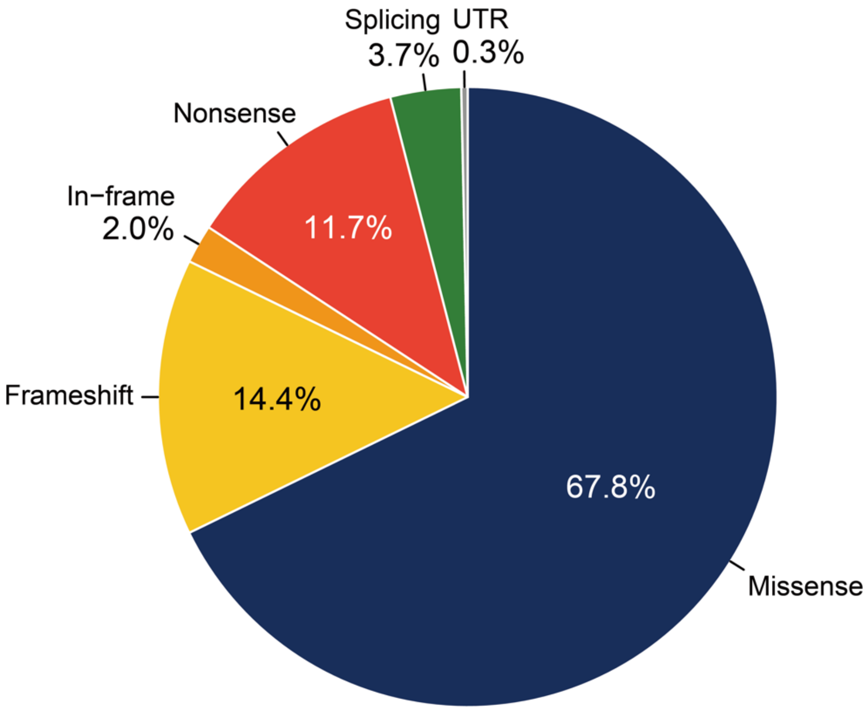


**Figure S5. Pathogenic variants categorized by function (*n* = 298 variants).**

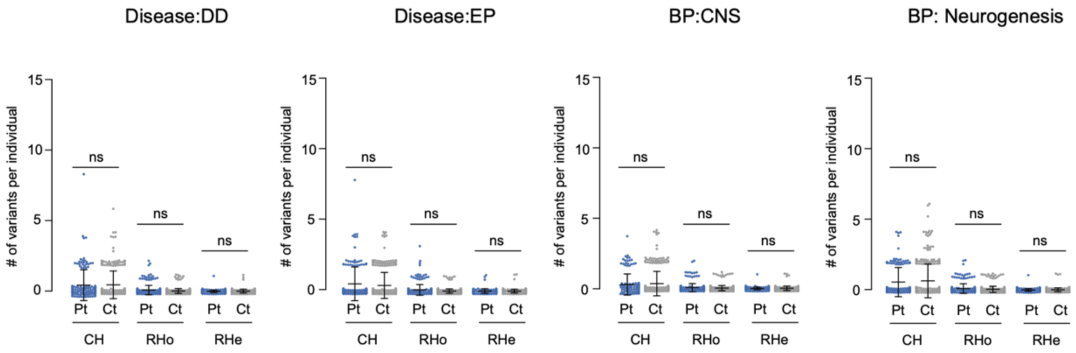


**Figure S6. No difference in the number of recessive variants for neurodevelopment-related gene sets between patients and controls.** All variants from genes from Disease:Developmental delay (DD), Disease:Epilepsy (EP), GO biological process (BP): central nervous system development (CNS) and GO BP:Neurogenesis.

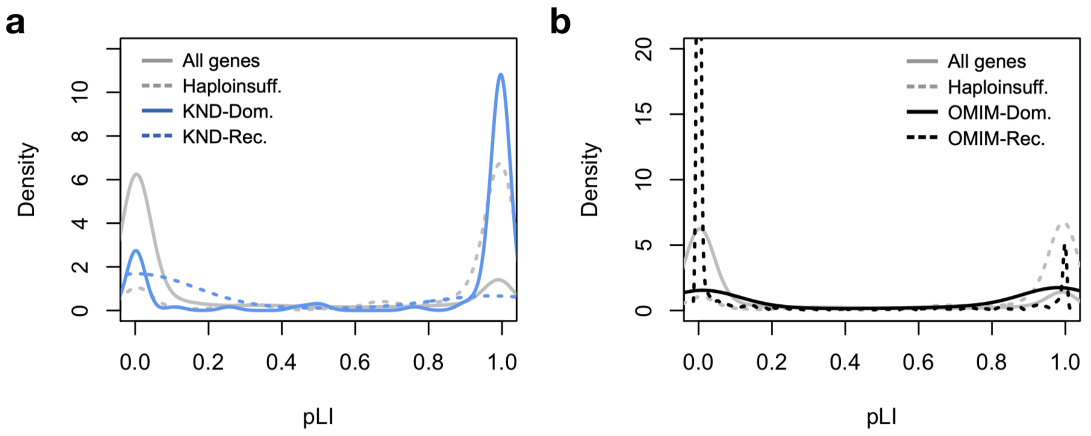


**Figure S7. Distribution of pLI values.**

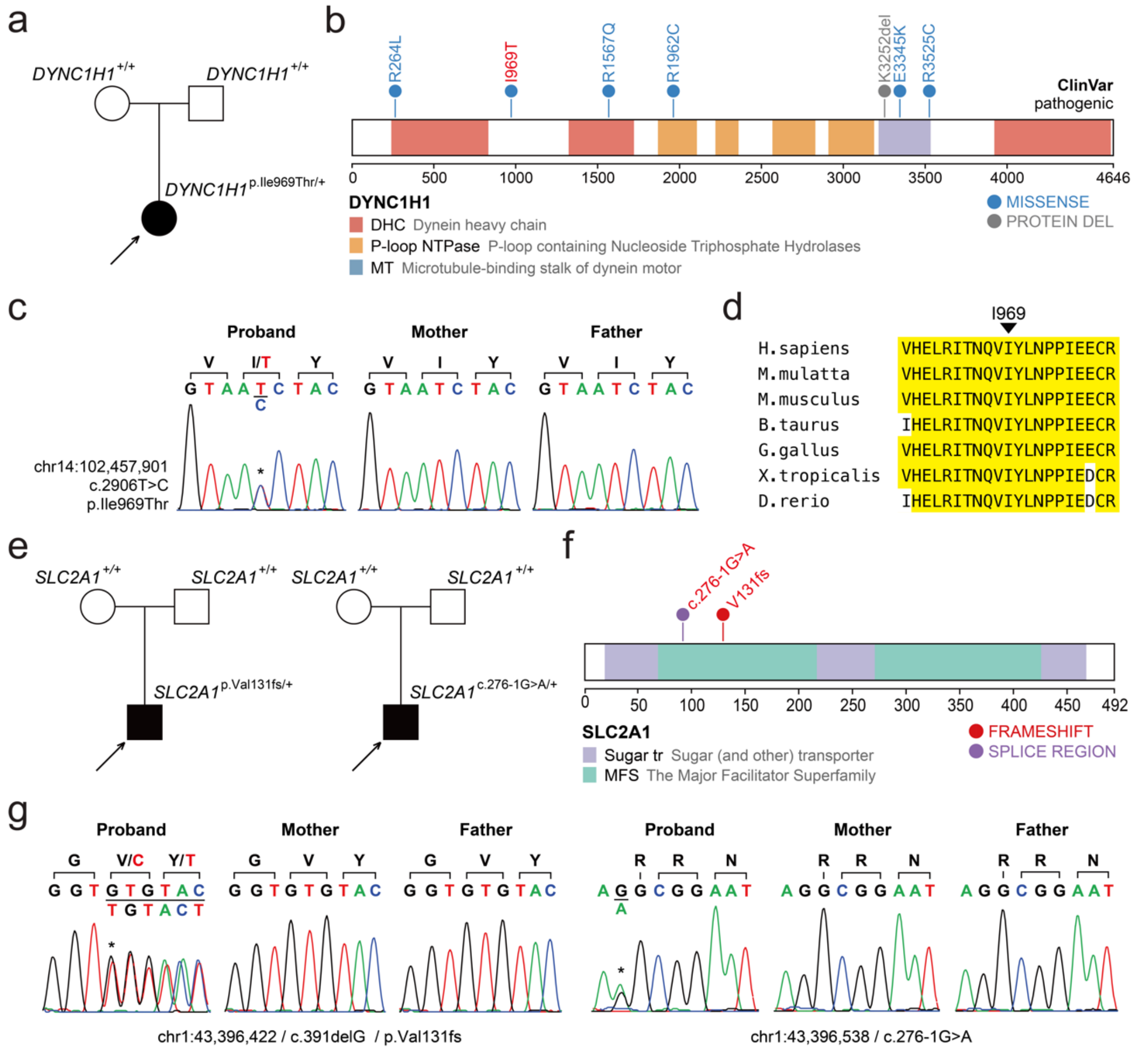


**Figure S8. Presentation and validation of cases that WES-based analysis altered clinical courses**. (a) A *de novo* variant in *DYNC1H1* identified from a patient with delayed development and hypotonia. (b) Pathogenic variants from ClinVar and domains of DYNC1H1 are displayed. A variant discovered in this study is shown in red. (c) Sanger traces validating the p.Ile969Thr variant. (d) Evolutionary conservation of the Ile969 residue across orthologs from major vertebrate species. (e) Two patients with dystonia and delayed motor development harbored LoF *de novo* variants in *SLC2A1*. (f) The variants found in two patients and domains of SLC2A1 are displayed. (g) Sanger traces validating the *de novo* variants.

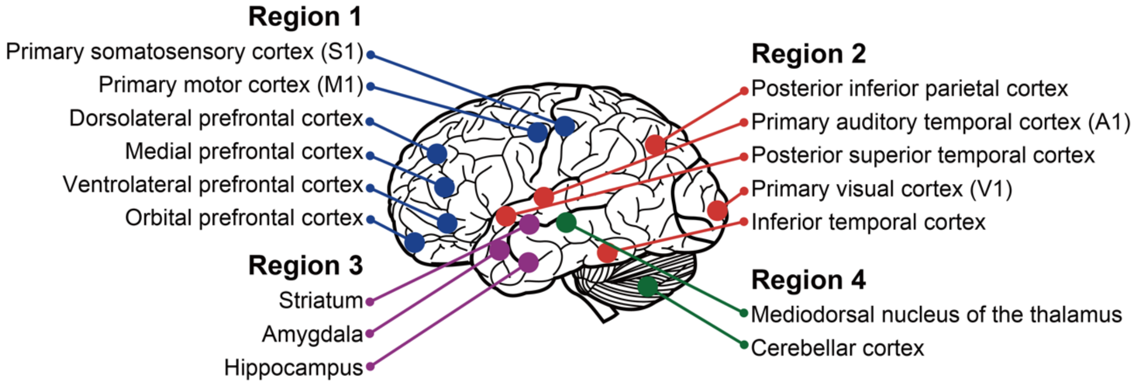


**Figure S9. Anatomical components of each brain regions that were used for novel gene network analysis.**

**Supplementary Table S1. List of copy number variations discovered in this study.**

| No. | Simple Diagnosis | CNV interval (Mb, hg19) | CNV length | Del/Dup | inheritance | No. of genes* | Notable gene(s) in the interval | Ref |
|---|---|---|---|---|---|---|---|---|
| **1** | Rett syndrome-like | chr1:0.6-3.1 | 2.6 Mb | Del | de novo | 60 | *MMP23B, GABRD, SKI, PRDM16* | [8] |
| **2** | Rett syndrome-like | chr1:107.0-113.1 | 6.1 Mb | Del | de novo | 69 | *WNT2B, NTNG1* | [9] |
| **3** | Hereditary spastic paraplegia | chr2:32.3-32.5 | 165.5 kb | Del | Inherited | 3 | *SPAST* | [10] |
| **4** | Epileptic encephalopathy | chr2:171.2-175.1 | 3.8 Mb | Del | de novo | 18 | *GAD1* | [11] |
| **5** | Unknown encephalopathy | chr2:234.8-242.8 | 8.0 Mb | Del | de novo | 61 | *HDAC4, PRLH, PER2, TWIST2, CAPN10, KIF1A, FARP2, D2HGDH, PDCD1* | [12, 13] |
| **6** | Rett syndrome-like | chr3:9.0-13.0 | 4.0 Mb | Del | de novo | 47 | *SETD5, SLC6A1, SLC6A11* | [14, 15] |
| **7** | Developmental delay with facial dysmorphism | chr3:44.2-48.0 | 3.9 Mb | Del | de novo | 55 | *SETD2, CSPG5, PTH1R, SMARCC1* | [16] |

| | | | | | | | | |
|---|---|---|---|---|---|---|---|---|
| **8** | Rett syndrome-like | chr4:0.0-2.8 | 2.8 Mb | Del | de novo | 42 | *PIGG, CPLX2, FGFRL1, CTBP1, SLBP, LETM1* | [17] |
| **9** | Congenital hypotonia | chr5:139.0-139.6 | 567.7 kb | Del | de novo | 7 | *NRG2, PURA* | [18] |
| **10** | Epileptic encephalopathy | chr7:119.7-119.9 | 171.3 kb | Dup | de novo | 0 | *KCND2* upstream | [19] |
| **11** | Epileptic encephalopathy | chr9:0-47.3 | 47.3 Mb | Dup | de novo | 202 | n.d. | [20] |
| **12** | Developmental delay with facial dysmorphism | chr9:139.6-141.1 | 1.6 Mb | Del | de novo | 56 | *EHMT1* | [21] |
| **13** | Developmental delay with facial dysmorphism | chr12:53.6-54.1 | 537.9 kb | Dup | de novo | 18 | *AAAS, AMHR2, RARG* | [22] |
| **14** | Developmental delay, Joubert syndrome | chr15:20.3-23.7 | 3.4 Mb | Del | de novo | 11 | *TUBGCP5, NIPA1, NIPA2, CYFIP1* | [23] |
| **15** | Intellectual disability, facial dysmorphism | chr16:29.6-30.2 | 635.4 kb | Del | de novo | 26 | *MVP, CDIPT1, SEZ6L2, ASPHD1* | [24] |
| **16** | Global Developmental delay | chr18:52.5-53.3 | 787.2 kb | Del | de novo | 3 | *TCF4* | [25] |
| **17** | Hereditary spastic paraplegia | chr22:21.1-21.6 | 508.7 kb | Del | de novo | 10 | *CRKL* | [26] |

| | | | | | | | | |
|---|---|---|---|---|---|---|---|---|
| **18** | Unknown<br>severe retardation | chr22:42.8-<br>51.3 | 8.6 Mb | Del | de novo | 81 | *SHANK3, IGF1* | [27] |
| **19** | Developmental delay,<br>microcephaly | chrX::41.4-<br>41.7 | 299.5 kb | Del | de novo | 3 | *CASK* | [28] |
| **20** | Unknown<br>encephalopathy | chrX::100.1-<br>107.3 | 7.2 Mb | Dup | hemizygous | 76 | *PLP1* | [29] |
| **21** | Neurodegeneration<br>with severe motor<br>developmental arrest | chrX:102.5-<br>103.2 | 692.2 kb | Dup | de novo | 15 | *PLP1* | [30] |
| **22** | Epileptic encephalopathy | chrX:152.8-<br>153.4 | 620.3 kb | Dup | de novo | 21 | *MECP2* | [31] |
| **23** | Hereditary spastic<br>paraplegia | chrX:153.6-<br>153.8 | 203.2 kb | Dup | hemizygous | 15 | *RPL10, ATP6P1, GDI1,<br>IKBKG* | [32, 33] |

* Based on NCBI RefSeq database. Only protein-coding genes were counted.

**Table S2. List of known neurologic disorder associated genes**

| Inheritance pattern | Gene | No. |
| --- | --- | --- |
| Autosomal-dominant | *ACOX1, ACTB, ACTG1, ADCY5, ANKRD11, ANO3, ARID1B(3), ARID2, ASXL3(4), ATL1(4), ATP1A3, BICD2, BPTF(2), C19orf12, CHD3, CHD7, CLTC, COL1A1(2), CTNNB1(5), DHX30, DNM1(2), DNM1L, DYNC1H1(2), EHMT1, FOXG1(2), FOXP1, GABBR2, GABRB1, GFAP, GNAO1(5), GNB1, GRIA2, GRIN1(3), GRIN2A, GRIN2B(3), HECW2, ITPR1(3), KAT6A(2), KCNC1, KCNQ2, KIF1A(3), KMT2A, LRP5, MED13L, MRAS, MYH7, NALCN(2), NFIX, NFKB2, NIPBL, NRAS, PIK3CA(3), PMP22, POGZ, PPP2R5D(2), PPP3CA, PTEN, PURA(2), RAB11B, RARB(3), REEP1(2), RHOBTB2(2), SATB2, SCN1A, SCN2A(3), SCN4A, SETD2, SFTPC, SLC2A1(2), SLC6A1, SLC6A5, SMARCB1, SOX10, SOX5, SPAST, SPTBN2, STAT3, STXBP12(2), SYNGAP1, TCF4(3), TUBB3, TUBB4A(4), UBE3A, WASHC5, WDFY3, ZEB2* | 86 |
| Autosomal-recessive | *ABAT, ALDH7A1, APTX, ARSA(2), ASAH1, ATAD3A, ATP6V0A2, BRAT1(2), CC2D2A, CLN6(2), CNTNAP1, CWF19L1, CYP7B1, DEGS1, ECHS1(2), EIF2B2, EIF2B3, ERCC5, FIG4, GCDH, GDAP1, GLB1(2), GMPPB(2), HADHA, HSD17B4, IARS2(2), IGHMBP2, KIAA1109, KIF1A,* | 55 |

| | | |
|---|---|---|
| | *KLHL40, LONP1(2), MICU1, NARS2(2), NDUFAF6(2), NDUFS1, NDUFV1, PEX5, PIGN, PLA2G6(2), PMM2, PNPT1(2), POLR1C, PPT1, PRUNE1, RYR1, SACS(2), SLC19A3(2), SLC25A15, ST3GAL5, SURF1, THOC6, TMEM173, TUBA8, WDR62, WDR81* | |
| X-linked | *ATRX(3), CASK(2), CDKL5(3), EDA, EIF2S3, GRIA3(2), HDAC8, HPRT1, IQSEC2(2), MECP2(3), MED12, NAA10, NSDHL, OPHN1(2), PAK3, PCDH19, PDHA1(2), PHF6, PLP1, SLC16A2, SMC1A(2), USP9X, ZC4H2(4)* | 23 |

( ): Number of patients (more than one patient only)

8. Jordan VK, Zaveri HP, Scott DA. 1p36 deletion syndrome: An update. Appl Clin

Genet. 2015;8:189–200.

8.
16. Lovrecic L, Bertok S, Žerjav Tanšek M. A New Case of an Extremely Rare 3p21.31 Interstitial Deletion. Mol Syndromol. 2016;7:93–8.
17. Battaglia A, Carey JC, South ST. Wolf-Hirschhorn syndrome: A review and update. Am J Med Genet Part C Semin Med Genet. 2015;169:216–23.
18. Hosoki K, Ohta T, Natsume J, Imai S, Okumura A, Matsui T, et al. Clinical phenotype and candidate genes for the 5q31.3 microdeletion syndrome. Am J Med Genet Part A. 2012;158 A:1891–6.
19. Zhao J, Noon SE, Krantz ID, Wu Y. A de novo interstitial deletion of 7q31.2q31.31 identified in a girl with developmental delay and hearing loss. Am J Med Genet Part C Semin Med Genet. 2016;172:102–8.
20. Abu-Amero KK, Hellani AM, Salih M a, Seidahmed MZ, Elmalik TS, Zidan G, et al. A de novo marker chromosome derived from 9p in a patient with 9p partial duplication syndrome and autism features: genotype-phenotype correlation. BMC Med Genet. 2010;11:135. doi:10.1186/1471-2350-11-135.
21. Kleefstra T, de Leeuw N. Kleefstra Syndrome. 2010 Oct 5 [Updated 2019 Mar 21]. In: Adam MP, Ardinger HH, Pagon RA, et al., editors. GeneReviews® [Internet]. Seattle (WA): University of Washington, Seattle; 1993-2019.
22. Jonsson DI, Ludvigsson P, Aradhya S, Sigurdardottir S, Steinarsdottir M, Hauksdottir H, et al. A de novo 1.13 Mb microdeletion in 12q13.13 associated with congenital distal arthrogryposis, intellectual disability and mild dysmorphism. Eur J Med Genet. 2012;55:437–40. doi:10.1016/j.ejmg.2012.03.001.
23. Doornbos M, Sikkema-Raddatz B, Ruijvenkamp CAL, Dijkhuizen T, Bijlsma EK, Gijsbers ACJ, et al. Nine patients with a microdeletion 15q11.2 between breakpoints 1 and 2 of the Prader-Willi critical region, possibly associated with behavioural

disturbances. Eur J Med Genet. 2009;52:108–15. doi:10.1016/j.ejmg.2009.03.010.

31. Van Esch H, Bauters M, Ignatius J, Jansen M, Raynaud M, Hollanders K, et al.

Duplication of the MECP2 region is a frequent cause of severe mental retardation and progressive neurological symptoms in males. Am J Hum Genet. 2005;77:442–53. doi:10.1086/444549.
32. Vandewalle J, Van Esch H, Govaerts K, Verbeeck J, Zweier C, Madrigal I, et al. Dosage-Dependent Severity of the Phenotype in Patients with Mental Retardation Due to a Recurrent Copy-Number Gain at Xq28 Mediated by an Unusual Recombination. Am J Hum Genet. 2009;85:809–22.
33. Fusco F, D'Urso M, Miano MG, Ursini MV. The LCR at the IKBKG Locus Is Prone to Recombine. Am J Hum Genet. 2010;86:650–2.
